# Targeting the Src pathway enhances the efficacy of selective FGFR inhibitors in cancers with FGFR3 alterations

**DOI:** 10.1101/2020.04.04.025544

**Authors:** Nadia Carvalho Lima, Eliza Atkinson, Tom D Bunney, Matilda Katan, Paul H. Huang

**Author notes:** Correspondence to: Paul H Huang, Division of Molecular Pathology, Institute of Cancer Research, 237 Fulham Road, London SW3 6JB, United Kingdom.

## Abstract

Selective FGFR inhibitors such as infigratinib (BGJ398) and erdafitinib (JNJ-42756493) have been evaluated in clinical trials for cancers with FGFR3 molecular alterations, particularly in urothelial carcinoma patients. However, a substantial proportion of these patients (up to 50%) display intrinsic resistance to these drugs and receive minimal clinical benefit. There is thus an unmet need for alternative therapeutic strategies to overcome primary resistance to selective FGFR inhibitors. In this study, we demonstrate that cells expressing cancer-associated activating FGFR3 mutants and the FGFR3-TACC3 fusion showed primary resistance to infigratinib in long-term colony formation assays in both NIH-3T3 and urothelial carcinoma models. We find that expression of these FGFR3 molecular alterations resulted in elevated constitutive Src activation compared to wildtype FGFR3 and that cells co-opted this pathway as a means to achieve intrinsic resistance to infigratinib. Targeting the Src pathway with low doses of the kinase inhibitor dasatinib synergistically sensitized multiple urothelial carcinoma lines harbouring endogenous FGFR3 alterations to infigratinib. Our preclinical data provides evidence that supports the use of dasatinib in combination with selective FGFR inhibitors as a means to overcome intrinsic drug resistance in the salvage therapy setting in cancer patients with FGFR3 molecular alterations.

## Introduction

Activating molecular alterations in the Fibroblast Growth Factor Receptor 3 (*FGFR3*) gene have been identified through large-scale next generation genomic sequencing efforts across a range of tumour types including urothelial carcinoma, lung squamous cell carcinoma, glioblastoma and myeloma (1-4). These molecular alterations primarily comprise of point mutations that span across the gene (e.g. S249C, R248C and K652E) and gene fusions (e.g. FGFR3-TACC3 fusion). The cancer type with the highest proportion of FGFR3 alterations is urothelial carcinoma with a frequency of 60%-80% in the non-muscle-invasive form and 15-20% in the muscle-invasive form harbouring these molecular aberrations (5-8). The most common FGFR3 point mutation found across cancer types is S249C in which the extracellular domain cysteine substitution leads to the formation of a disulphide bond between receptor monomers resulting in ligand-independent constitutive dimerization and activation (2, 9-11). In contrast, the FGFR3-TACC3 fusion has been described at low frequencies (2-8%) in urothelial cancers and glioblastoma and mediates it oncogenic activity through constitutive dimerization of FGFR3 driven by TACC3 (1, 2, 4, 12, 13).

The identification of these FGFR3 aberrations in cancer has led to the clinical evaluation of selective Fibroblast Growth Factor Receptor (FGFR) small molecule tyrosine kinase inhibitors (TKIs) in this molecularly defined subset of patients. The most advanced candidates include erdafitinib (JNJ-42756493) and infigratinib (BGJ398). Pal et al., reported response rates of 25.4% in the evaluation of infigratinib in 67 patients with previously treated advanced urothelial carcinoma with FGFR3 alterations (14). Similarly, the BLC2001 phase II single-arm trial of erdafitinib in FGFR-altered urothelial carcinomas showed clinical responses in 40% of patients (15). This result has led to the accelerated approval of erdafitinib in 2019 by the US Food and Drug Administration (FDA) for locally advanced or metastatic urothelial carcinomas with FGFR2 and FGFR3 alterations. Despite these promising results, the BLC2001 trial showed that only 49% and 16% of patients with point FGFR mutations and fusions, respectively, achieved disease response with erdafitinib. Similarly, Pal et al., reported that only 42.9% of patients with documented FGFR3 activating point mutations responded to infigratinib (14, 15). These results indicate that notwithstanding the presence of a FGFR3 molecular alteration, there remains a significant proportion of patients that harbour intrinsic resistance to selective FGFR TKIs and do not respond to these agents. For these patients, there are limited therapeutic alternatives available and there is thus an urgent unmet need to identify effective ways to overcome primary resistance.

In this study, we show that cells expressing a subset of activating FGFR3 mutants and the FGFR3-TACC3 fusion are intrinsically resistant to selective FGFR TKI inhibition in long-term assays. We further demonstrate that expression of these FGFR3 mutants and FGFR3-TACC3 results in elevated constitutive Src activation levels compared to wildtype (WT) FGFR3, which can be exploited for therapy with the multi-target TKI dasatinib. We find that targeting the Src pathway with low dose dasatinib sensitizes urothelial carcinoma cell lines harbouring endogenous FGFR3 molecular alterations to selective FGFR TKIs including infigratinib. Our preclinical data suggests that patients with FGFR3 molecular alterations may benefit from treatment with FGFR and Src inhibitors in combination, which can overcome intrinsic resistance associated with selective FGFR TKI monotherapy.

## Results

### FGFR3-TACC3 gene fusion and activating FGFR3 point mutants are intrinsically resistant to infigratinib in the NIH-3T3 cell line model

We first evaluated the effects of infigratinib in NIH-3T3 cells expressing a selection of the most prevalent activating FGFR3 molecular alterations in cancer. These included the most common FGFR3 fusion (FGFR3-TACC3) and extracellular domain (S249C) and kinase domain (K652E) point mutations. In addition, we examined N542K, which has previously been described as a congenital mutation in hypochondroplasia (16), due to its reported high kinase activity in *in vitro* assays (17). Expression of the FGFR3-TACC3 gene fusion and FGFR3 mutations S249C, N542K and K652E led to constitutive phosphorylation of Src when compared to the WT FGFR3 control with no differences observed in Erk1/2 and AKT phosphorylation levels (Figure 1A). Consistent with previous reports, the transforming potential of these FGFR3 aberrations was demonstrated by an increase in the ability to grow under anchorage-independent conditions versus WT FGFR3 (Figure 1B-C) (17-19). Short-term (72 hours) cell viability experiments finds that as previously shown (17, 20-22), cells expressing FGFR3-TACC3 and the three FGFR3 mutants were sensitive to treatment with infigratinib with IC_50_ values in the nanomolar range (Figure 1D-E). It has been demonstrated that FGFR3 inhibition with FGFR inhibitors leads to selective downregulation of the Erk1/2 pathway (22-24). We show that unlike the WT FGFR3 cells, treatment with infigratinib decreased phosphoErk1/2 levels in the three mutant FGFR3 lines and abolished Erk1/2 phosphorylation in a dose-dependent manner in the FGFR3-TACC3 cells, confirming that this pathway is a direct downstream target of the FGFR3 mutants and FGFR3-TACC3 fusion. Interestingly, although constitutive activation of Src was associated with the expression of the FGFR3 mutants and FGFR3-TACC3 fusion in the NIH-3T3 cells (Figure 1A), the phosphorylation levels of Src were not downregulated upon FGFR3 kinase inhibition by infigratinib (Figure 1F). Further assessment of the efficacy of infigratinib in long-term (2 weeks) colony formation assays finds that despite the potent effects of this drug in the FGFR3 mutant and fusion expressing cells in short-term cell viability assays; there was an unexpected observation of a high number of resistant colonies persisting after 2 weeks of drug treatment (Figure 1G-H). At 1µM drug treatment, there was no significant decrease in colony formation in any of the FGFR3 point mutant expressing cells, with only a small decrease (∼20%) in the FGFR3-TACC3 expressing cells (Figure 1H). Importantly, these intrinsic resistant colonies persist even at very high infigratinib doses of 2uM which models the primary resistance observed in a subset of patients in clinical trials.

**Figure 1.**
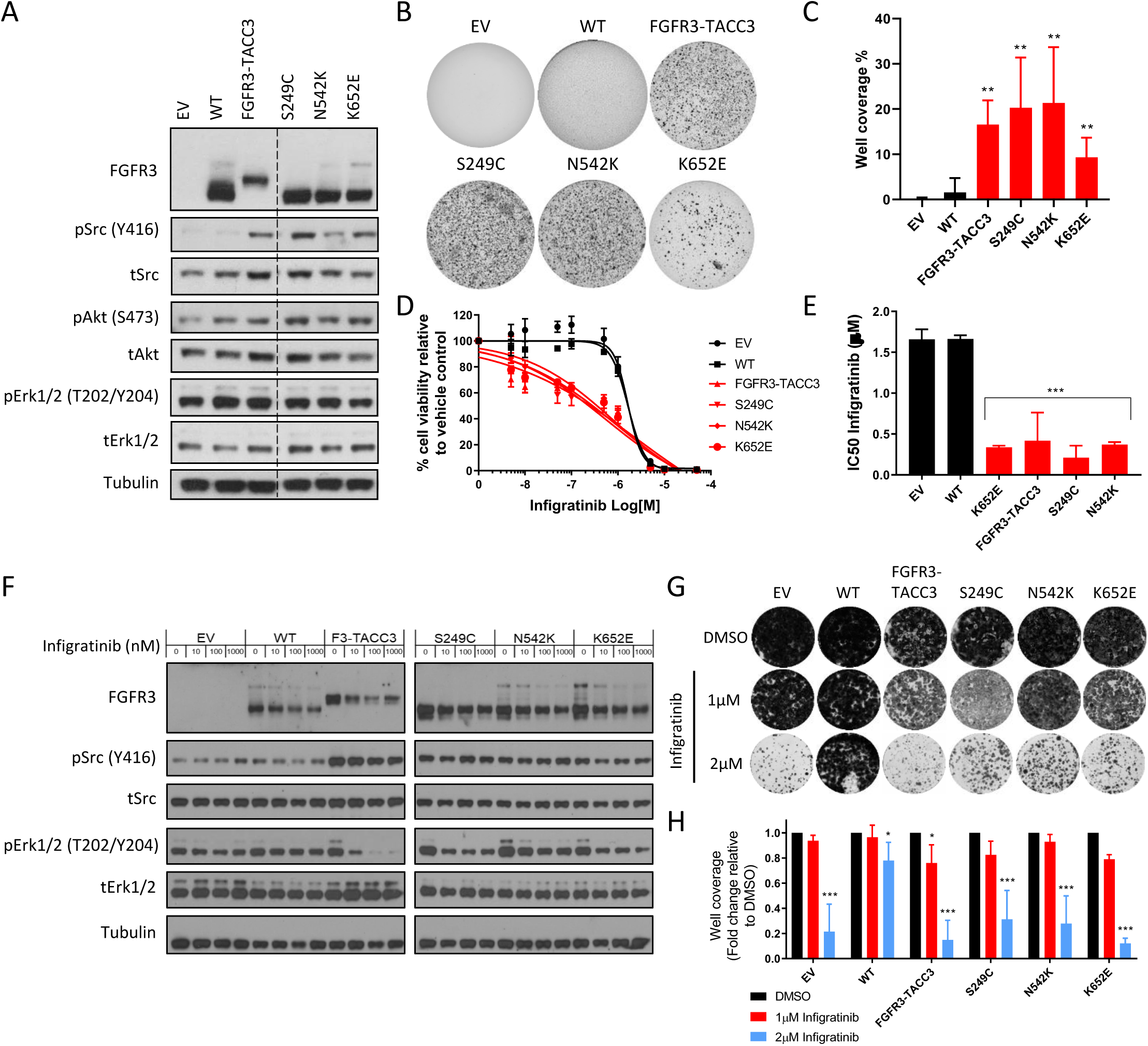
Cells bearing FGFR3 molecular alterations harbour intrinsic resistance to infigratinib. (A) Immunoblot of downstream cellular signalling pathways upon expression of FGFR constructs in NIH3T3 cells is shown. (B) Representative images of the panel of NIH-3T3 cells in anchorage-independent soft agar assays. (C) Bar plots showing the quantification of well coverage of soft agar assay in panel B (n=6). Statistical analysis of FGFR3 mutants and fusion versus WT FGFR3 was performed by one-way ANOVA with Dunnett’s post-hoc multiple comparison adjusted p-value (**p < 0.01). (D) Dose-response curves of the panel of NIH-3T3 cell lines upon treatment with infigratinib for 72 hours. Cell viability is normalised to DMSO control treatment (n=3). (E) Bar plots showing IC_50_ values of infigratinib calculated from dose response curves in (D). Statistical analysis of FGFR3 mutations and fusion versus WT FGFR3 was performed by one-way ANOVA with Dunnett’s post-hoc multiple comparison adjusted p-value (***p < 0.001). (F) Immunoblot of Erk1/2 and Src phosphorylation levels in NIH-3T3 cells treated with infigratinib at the indicated doses for 6 hours is shown. (G) Representative images of long term colony formation assay in the NIH-3T3 cell line panel upon treatment with infigratinib or DMSO control at the indicated doses for 2 weeks. (I) Bar plots showing the quantification of well coverage of the colony formation assay in panel G. Data for each cell line is normalised to DMSO control treatment (n=3). Statistical analysis of drug treatment versus DMSO was performed by paired two-way ANOVA with Dunnett’s post-hoc multiple comparison adjusted p-value (*p<0.05, ***p<0.001). Data presented for (C), (D), (E) and (H) represent mean ± SD. EV – empty vector control, WT – wildtype FGFR3, F3-TACC3 – FGFR3-TACC3.

### FGFR3-TACC3 and S249C expression confers resistance to dasatinib

To interrogate the key signalling dependencies in the panel of FGFR3 expressing NIH-3T3 cells, a targeted small molecule inhibitor screen was undertaken. This screen was comprised of 32 small molecule inhibitors that target major kinase and non-kinase oncogenic signalling pathways in cells. These inhibitors include broad-spectrum kinase inhibitors such as imatinib, dasatinib and foretinib as well as selective kinase inhibitors such as infigratinib (FGFR), binimetinib (MEK), AZD5363 (AKT), NVP-BEZ235 (PI3K/mTOR) and MK8776 (CHK1). The screen also has a small number of non-kinase inhibitors including NVP-AUY922 (HSP90), GSK126 (enhancer of zeste homolog 2 (EZH2)) and JQ1 (bromodomain and extra-terminal (BET)) (See Table S1 for list of compounds used in the screen and key targets).

As a positive control for the assay, we show that as expected, FGFR3 point mutant and fusion expressing cells were sensitive to both multi-target (ponatinib, foretinib, lenvatinib, cediranib) and selective (infigratinib and AZD4546) FGFR TKIs (Figure 2A). Interestingly, the expression of some FGFR3 molecular alterations conferred a survival advantage (of >1.2 fold) to a small number of compounds compared to WT FGFR3. These included BEZ235 (N542K and N652E), JQ1 (FGFR-TACC3, S249C and N652E), MK8776 (S249C) and dasatinib (FGFR3-TACC3 and S249C). Given that the expression of FGFR3 molecular alterations led to the constitutive activation of Src (Figure 1A), dasatinib, a broad-spectrum TKI that potently inhibits Src as one of its targets (25, 26), was taken forward for further investigation.

**Figure 2.**
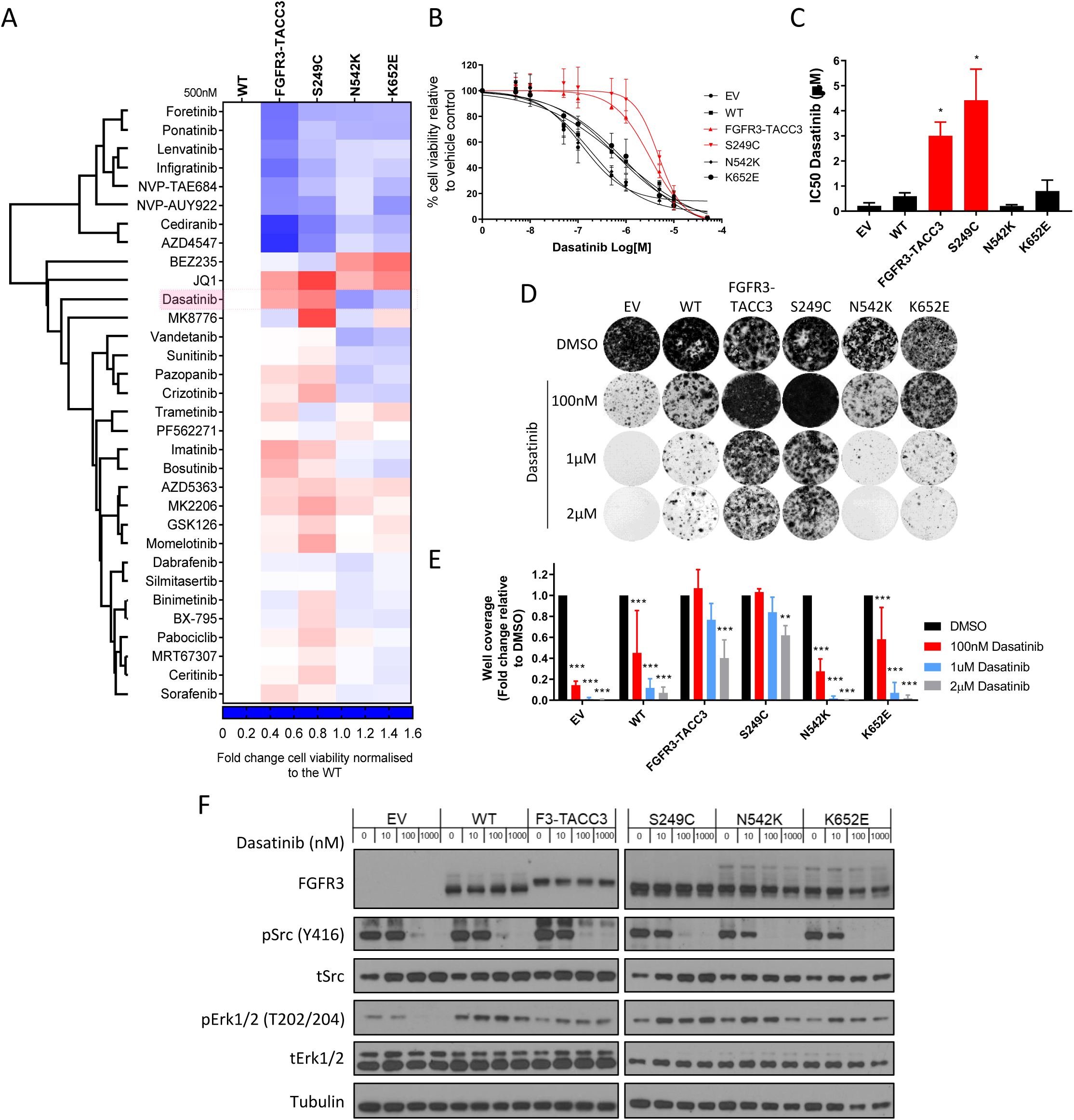
Cells expressing FGFR3-TACC3 and FGFR3 S249C mutant are resistant to dasatinib as a single agent. (A) Heatmap depicting one-way hierarchical clustering of cell viability data in targeted drug screen. FGFR3 expressing NIH-3T3 cells were seeded in 96 well places and viability measured using CTG assay following 72 hours treatment with small molecular inhibitors at 500nM (50nM for NVP-AUY=922). Cell viability data is normalised to DMSO control for each cell line and represented in heatmap relative to WT FGFR3 cells (n=3). (B) Dose-response curves of the panel of NIH-3T3 cell lines upon treatment with dasatinib for 72 hours. Cell viability is normalised to DMSO control treatment (n=3). (C) Bar plots showing IC_50_ values of dasatinib calculated from dose response curves in (B). Statistical analysis of FGFR3 mutations and fusion versus WT FGFR3 was performed by one-way ANOVA with Dunnett’s post-hoc multiple comparison adjusted p-value (*p < 0.05). (D) Representative images of long term colony formation assay in the NIH-3T3 cell line panel upon treatment with dasatinib or DMSO control at the indicated doses for 2 weeks. (E) Bar plots showing the quantification of well coverage of the colony formation assay in panel D. Data for each cell line is normalised to DMSO control treatment (n=3). Statistical analysis of drug treatment versus DMSO was performed by two-way ANOVA with Dunnett’s post-hoc multiple comparison adjusted p-value (**p<0.01, ***p<0.001). (F) Immunoblot of Erk1/2 and Src phosphorylation levels in NIH-3T3 cells treated with infigratinib at the indicated doses for 6 hours is shown. Data presented for (B), (C) and (E) represent mean ± SD. EV – empty vector control, WT – wildtype FGFR3, F3-TACC3 – FGFR3-TACC3.

Full dose response assessment of dasatinib in short-term viability assays showed that FGFR3-TACC3 and S249C conferred resistance to this drug with an IC_50_ of 3-4µM compared to the WT FGFR3 and the N542K and K652E mutations (<1µM) (Figure 2B-C). These results were recapitulated in the long-term colony formation assays with the FGFR3-TACC3 and S249C expressing cells showing high levels of resistant colonies in the presence of dasatinib compared to the other cell lines in the panel (Figure 2D-E). Evaluation of downstream signalling pathways by immunoblotting confirmed that dasatinib treatment led to a dose dependent decrease in Src phosphorylation levels across the cell line panel (Figure 2F).

We hypothesized that the constitutive dimerisation in these FGFR3 molecular alterations may be responsible for the observed resistance to dasatinib. Since disruption of the FGFR3-TACC3 fusion through genetic means is non-trivial, we focused our efforts on the S249C point mutation. It has previously been demonstrated that the cysteine substitution in S249 within the extracellular domain of FGFR3 leads to constitutive receptor dimerisation and activation through the formation of an intermolecular disulphide bond (2, 9-11). To investigate the role of this cysteine residue in driving resistance to dasatinib, we engineered NIH3T3 cell lines expressing an FGFR3 mutant where the cysteine has been substituted to an alanine residue (S249A). Evaluation in non-reducing PAGE showed that the alanine substitution led to the loss of constitutive FGFR3 dimers both in the presence and absence of infigratinib (Figure S1A). While cells expressing S249C were resistant to dasatinib in both short-term viability assays and long-term colony formation assays, the substitution to alanine resulted in pronounced sensitivity to this drug with levels comparable to WT FGFR3 expressing cells (Figure S1B-D). Similar results were seen in cells engineered to express another common FGFR3 cysteine mutant found in urothelial carcinoma (R248C) and its glycine substituted mutant (R248G) (Figure S1E-F). Taken together, our data demonstrates that while treatment with dasatinib resulted in a decrease in Src phosphorylation levels in all FGFR3 expressing cell lines examined, the expression of FGFR3-TACC3 and S249C unexpectedly confers resistance to this drug. Furthermore, we show that in two of the most common FGFR3 extracellular cysteine mutants in urothelial carcinoma (S249C and R248C), the constitutive receptor dimerisation mediated by the cysteine residue is necessary for driving dasatinib resistance.

### Dasatinib sensitizes NIH-3T3 cells with FGFR3 molecular alterations to infigratinib treatment

The use of combination therapy as a means to suppress multiple signalling pathways to overcome drug resistance is a well-established concept in cancer biology (27, 28). Given that infigratinib and dasatinib act through parallel signalling pathways of Erk1/2 and Src respectively (Figure 1F and 2F), we posited that the addition of dasatinib may enhance the activity of infigratinib through the dampening of the Src pathway to overcome the intrinsic resistance observed in long-term colony formation assays (Figure 1G). Co-treatment of infigratinib with low dose (100-200nM) of dasatinib resulted in an almost complete elimination of the resistant colonies that were previously observed in the treatment with single agent infigratinib or dasatinib (Figure 3A-B). Notably, this effect appeared to be more potent in cells expressing FGFR3-TACC3 and the FGFR3 point mutants with some resistant colonies remaining in the WT FGFR3 expressing cells even at high drug doses (Figure 3B). This result was recapitulated in the short-term cell viability assays with the FGFR3-TACC3 and S249C expressing cells having a more pronounced sensitivity to the combination than WT FGFR3 expressing cells (Figure 3C).

**Figure 3.**
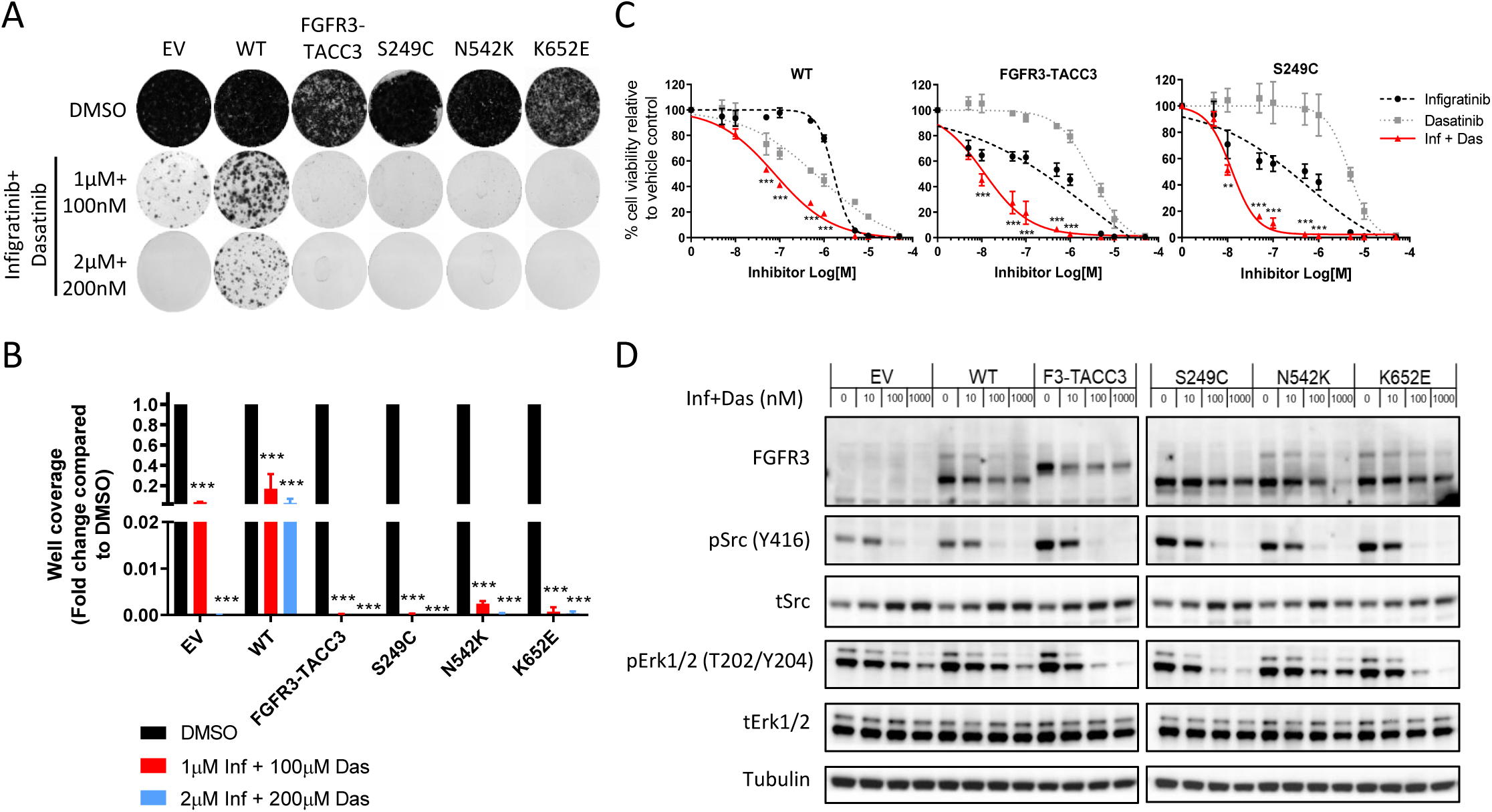
Combination of dasatinib and infigratinib overcomes intrinsic resistance. (A) Representative images of long term colony formation assay in the NIH-3T3 cell line panel upon treatment with a combination of dasatinib and infigratinib or DMSO control at the indicated doses for 2 weeks. (B) Bar plots showing the quantification of well coverage of the colony formation assay in panel A. Data for each cell line is normalised to DMSO control treatment (n=3). Statistical analysis of drug treatment versus DMSO was performed by two-way ANOVA with Dunnett’s post-hoc multiple comparison adjusted p-value (***p<0.001). (C) Dose-response curves of the WT FGFR3, FGFR3-TACC3 and S249C cell lines upon treatment with infigratinib and dasatinib either as single agents or in combination for 72 hours. For the combination arm, cells were treated with infigratinib and dasatinib in a 1:1 ratio with each drug at half the dose of the single agent. Cell viability is normalised to DMSO control treatment (n=3). Statistical analysis of the combination arm versus either single agent at each dose was evaluated using two-way ANOVA with Dunnett’s post-hoc multiple comparison adjusted p-value at each drug dose (**p<0.01, ***p<0.001). (D) Immunoblot of Erk1/2 and Src phosphorylation levels in NIH-3T3 cells treated with a combination of infigratinib and dasatinib in a 1:1 ratio for 6 hours, with the indicated concentration representing to total drug concentration used. Data presented for (B) and (C) represent mean ± SD. EV – empty vector control, WT – wildtype FGFR3, F3-TACC3 – FGFR3-TACC3, Inf – infigratinib, Das – dasatinib.

To evaluate the signalling consequence of combination therapy with infigratinib and dasatinib, an assessment of Erk1/2 and Src activation levels after treatment with both drugs was performed by immunoblotting. As expected, cells bearing FGFR3-TACC3, S294C, N542K and K652E showed a decrease in the phosphorylation levels of both Src and Erk1/2 upon treatment with the combination, which is the result of the independent effects of dasatinib and infigratinib on these pathways respectively as single agents (Figure 3D). In the S249C and K652E expressing cells, there was a more potent inhibition of phosphoErk1/2 in the drug combination compared to single agent treatment with infigratinib alone (Figure 1F), suggesting that in a subset of FGFR3 mutants, dasatinib may enhance the effects of infigratinib by increased suppression of the Erk1/2 pathway.

### Targeting the Src pathway enhances the efficacy of selective FGFR inhibitors in urothelial cancer cell lines with FGFR3 molecular alterations

We sought to establish if the drug responses observed in the NIH-3T3 model system is also present in cancer cell lines harbouring endogenous FGFR3 molecular alterations. To that end, we evaluated the effects of two structurally unrelated selective FGFR inhibitors (infigratinib and PD173074) and dasatinib as single agents or in combination in a panel of urothelial cancer cell lines. The panel included RT112M (FGFR3-TACC3) and 639V (R248C) as well as BFTC905 which expresses WT FGFR3 and was used as a control (29-31). Long-term colony formation assays show that in line with previous reports, RT112M was partially sensitive to infigratinib and PD173074 as single agents compared to BFTC905 and 639V (Figure 4A) (23). It should be noted that despite the comparatively higher sensitivity observed in the RT112M cells, there were still a large proportion of residual colonies upon drug treatment, indicating that a significant fraction of cells displayed intrinsic resistance to selective FGFR inhibitors. As observed in the NIH-3T3 panel, WT FGFR3 expressing BFTC905 cells were sensitive to dasatinib treatment as a single agent while RT112M and 639V cells with FGFR3 molecular alterations were partially resistant to this drug. The addition of dasatinib at low dose enhanced the efficacy of both selective FGFR inhibitors in the RT112M and 639V cells resulting in a decrease in colony formation in the combination arm compared to single agent FGFR inhibitor treatment (Figure 4A-B). We further evaluated if the dasatinib and infigratinib combination was synergistic in dose response cell viability assays (Figure 4C). Assessment of the combination index (CI) showed that the combination of dasatinib and infigratinib led to a synergistic decrease in the cell viability (CI <1) in the RT112M and 639V cells across all drug doses tested up to 1µM (Figure 4D).

**Figure 4.**
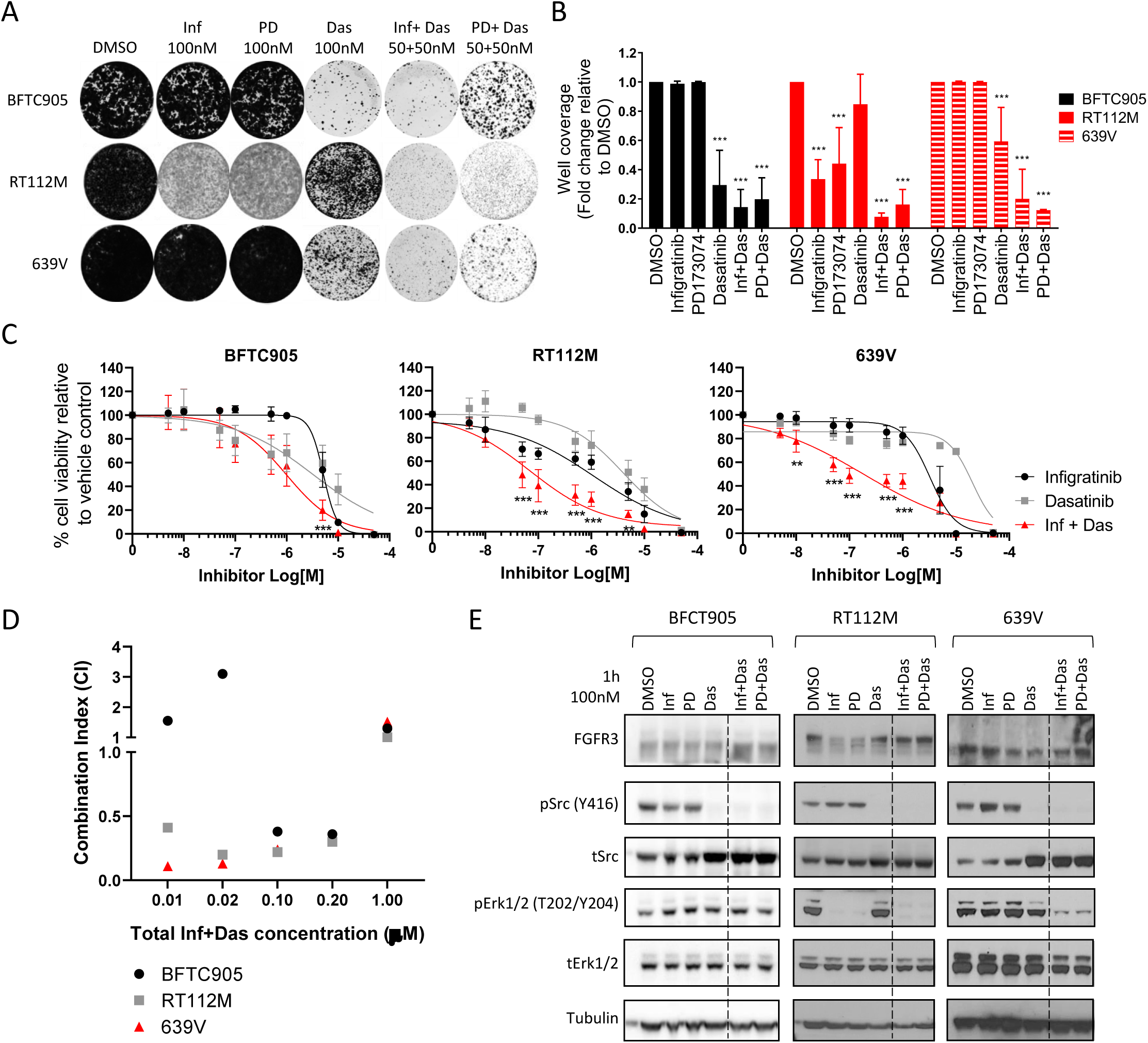
Targeting the Src pathway enhances the efficacy of selective FGFR inhibitors in urothelial cancer cell lines with FGFR3 molecular alterations. (A) Representative images of long term colony formation assay in urothelial cancer cell line panel upon treatment with infigratinib, PD173074, dasatinib as single agent or in combination as well as DMSO control at the indicated doses for 2 weeks. (B) Bar plots showing the quantification of well coverage of the colony formation assay in panel A. Data for each cell line is normalised to DMSO control treatment (n=3). Statistical analysis of drug treatment versus DMSO was performed by two-way ANOVA with Dunnett’s post-hoc multiple comparison adjusted p-value (***p<0.001). (C) Dose-response curves of the urothelial cancer cell lines upon treatment with infigratinib and dasatinib either as single agents or in combination for 72 hours. For the combination arm, cells were treated with infigratinib and dasatinib in a 1:1 ratio with each drug at half the dose of the single agent. Cell viability is normalised to DMSO control treatment (n=3). Statistical analysis of the combination arm versus either single agent at each dose was evaluated using two-way ANOVA with Dunnett’s post-hoc multiple comparison adjusted p-value at each drug dose (**p<0.01, ***p<0.001). (D) Combination index (CI) measurements for infigratinib and dasatinib in urothelial cancer cell lines based on dose response data in panel C. Synergy (CI < 1) in RT112M and 639V is shown across all doses tested up to 1µM. (E) Immunoblot of Erk1/2 and Src phosphorylation levels in urothelial cancer cells treated with single agent selective FGFR inhibitor or dasatinib and in combination for 1 hour. The combination arm was administered in a 1:1 ratio, with the indicated concentration representing the total drug concentration used. Data presented for (B) and (C) represent mean ± SD. Inf – infigratinib, PD – PD173074, Das – dasatinib.

To assess the signalling alterations associated with the observed phenotypes in the urothelial cancer cell lines upon drug treatment, immunoblotting of Erk1/2 and Src phosphorylation levels was undertaken. Infigratinib and PD173074 treatment did not affect phosphoErk1/2 levels in the BFTC905 and 639V cells but abolished phosphoErk1/2 levels in the RT112M cell line (Figure 4E). Dasatinib treatment suppressed Src phosphorylation in all three cell lines which was accompanied by an increase of total Src levels. As observed in the NIH3T3 cells, the combination of a selective FGFR inhibitor with dasatinib led to a reduction in the phosphorylation levels of both Erk1/2 and Src in the two cell lines harbouring FGFR molecular alterations, RT112M and 639V. The Epidermal Growth Factor Receptor (*EGFR*) has previously been shown to be a compensatory bypass mechanism which confers intrinsic resistance to PD173074 treatment in the RT112M cell line (23). Since Src is known to be regulated by EGFR, we set out to ascertain if the observed Src phosphorylation in the urothelial cancer cell lines is influenced by EGFR activity. Treatment of the cell line panel with the EGFR inhibitor gefitinib either as a single agent or in combination with the selective FGFR inhibitors did not lead to a reduction in Src phosphorylation levels (Figure S2A), confirming that Src is not an effector of EGFR activation but rather a parallel resistance pathway in urothelial cancer cell lines.

Given that dasatinib is a broad spectrum kinase inhibitor that blocks multiple kinases including Src (25, 26), we sought to demonstrate that Src is the primary target of this drug and the causative driver of the observed intrinsic resistance to selective FGFR inhibitors. Specifically, we evaluated the ability of the gatekeeper mutation in the avian Src gene (T338I), which blocks dasatinib binding to its kinase domain, to rescue the phenotypic effects of dasatinib (32). As a comparator, the WT Src gene and an empty vector (EV) control plasmid were used. Engineered urothelial cancer cell lines expressing each of the Src variants or the EV control were subjected to long-term colony formation assays in the presence of infigratinib and dasatinib as single agents or in combination. Expression of the gatekeeper mutant rescued the effects of dasatinib, conferring robust resistance to this drug in all three cell lines when compared to the WT Src and EV cells (Figure S2B). Collectively, these experiments provide evidence that low dose dasatinib is highly effective in overcoming intrinsic resistance to selective FGFR inhibitor treatment in urothelial cancer cell lines expressing FGFR3 molecular alterations and that dasatinib mediates this effect primarily through blockade of the Src pathway.

## Discussion

Despite some clinical success with selective FGFR inhibitors in cancers with FGFR3 molecular alterations, these therapies are not curative and there remains a substantial patient population that harbour intrinsic resistance to these drugs (14, 15). There is a need to consider new treatment strategies to tackle intrinsic drug resistance in order to improve long-term outcomes in patients. In this study, we use a combination of NIH-3T3 and urothelial cancer cells to demonstrate that cells expressing a subset of FGFR3 molecular alterations co-opt the Src pathway to promote intrinsic resistance to selective FGFR inhibitors. While cells expressing the FGFR3-TACC3 fusion and extracellular cysteine mutants were resistant to the Src inhibitor dasatinib and selective FGFR inhibitor infigratinib as single agents, the addition of low dose dasatinib profoundly sensitized these cells to selective FGFR inhibitors. Our data provides evidence that supports the use of this drug combination as a means to either overcome intrinsic drug resistance to FGFR inhibitors in the salvage therapy setting or as an upfront strategy to achieve durable drug responses in cancer patients with FGFR3 molecular alterations.

Preclinical and clinical studies have shed light on some of the mechanisms of resistance to selective FGFR inhibitors. These are broadly divided into on-target and bypass mechanisms. While on-target resistance mutations in the FGFR2 kinase domain (N649H/K, V564F, E656A, L617V, K641R and K659M) have been detected in circulating tumour DNA in the context of FGFR2 translocations in cholangiocarcinoma patients who have received infigratinib (33, 34), such mutations have yet to be identified in cancers with FGFR3 molecular alterations. Preclinical studies have shown that upregulation of the AKT pathway as well as ligand-mediated activation of ErbB2/3 are bypass signalling pathways in the context of acquired resistance to infigratinib in the RT112M cell line (35, 36). In the paradigm of intrinsic resistance, Herrera-Abreu et al., has shown that EGFR activation is an escape mechanism to PD173074 treatment and that a combination of gefitinib and PD173074 could overcome drug resistance in urothelial cancer lines harbouring FGFR3 molecular alterations (23). Our study defines Src as a new bypass pathway by which urothelial cancer cells achieve intrinsic resistance to selective FGFR inhibitors. We further demonstrate that in the urothelial cancer models examined, Src is not an effector of the EGFR-mediated mechanism previously described by Herrara-Abreu et al., but rather a parallel resistance pathway that is independent of EGFR signalling. To our knowledge, our report is the first description of the use of Src inhibitors such as dasatinib as a means to overcome intrinsic resistance to selective FGFR inhibitors in cancers with FGFR3 molecular alterations.

Dasatinib has been previously evaluated as a monotherapy in urothelial carcinoma. RT4 cells, which expresses a different form of the FGFR3-TACC3 fusion employed in this study, are sensitive to treatment with dasatinib both *in vitro* and *in vivo* (37). Interestingly, it has been shown in another study that genetic silencing of Src in RT4 cells promotes cancer cell migration and invasion (38). Dasatinib has been clinically evaluated in the neoadjuvant setting in muscle-invasive urothelial carcinoma (39). This phase II trial showed that while treatment with this drug led to an on-target decrease in the phosphorylation of Src, there was no clinical benefit as reflected by the lack significant changes in proliferation and apoptosis markers in pre- and post-treatment tumours (39). Our data shows that expression of FGFR3-TACC3 and S249C results in a paradoxical resistance to dasatinib despite the constitutive activation of the Src pathway, reinforcing the idea that this drug as a monotherapy is likely to be ineffective in cancers with these FGFR3 molecular alterations. One of the primary ways to address intrinsic drug resistance is the use of combination therapies that abrogate compensatory survival mechanisms (27, 40). We find that co-targeting FGFR3 and Src with low nanomolar concentrations of infigratinib and dasatinib respectively is highly effective in reducing cell viability in urothelial cancer cell lines. This drug combination simultaneously abrogates Erk1/2 and Src phosphorylation leading to durable efficacy in long term-colony formation assays, suggesting that simultaneous blockade of these parallel survival pathways is necessary to kill these cells. The use of Src inhibitors such as dasatinib to overcome resistance to kinase inhibitors has previously been investigated in the context of mutant EGFR lung cancer (41, 42). A phase I clinical trial of dasatinib in combination with the EGFR inhibitor erlotinib in advanced non-small-cell lung cancer showed that the combination is tolerable with disease control in a subset of patients (43). The clinical experience in lung cancer suggests that it is feasible to evaluate the efficacy of dasatinib in combination with selective FGFR inhibitors in patients with FGFR3 molecular alterations.

There remain several outstanding questions. For instance, the mechanisms by which Src blockade sensitizes cells to FGFR3 inhibition are poorly understood. Since expression of the FGFR3-TACC3 fusion and the substitution of extracellular cysteines in FGFR3 confers resistance to dasatinib as a single agent, it is tempting to speculate a mechanism whereby dasatinib treatment alone has no therapeutic effect, while the addition of a selective FGFR inhibitor induces a new dependency to the Src pathway leading to vulnerability to co-treatment with dasatinib. Such induced collateral sensitivity has been observed with other tyrosine kinase inhibitors (44, 45). There is also a need to investigate the efficacy of this combination beyond the small panel of FGFR3 molecular alterations examined in this study in order to establish the generality of these findings and expand its utility to patients harbouring other FGFR3 mutations. Furthermore, it remains unclear why some FGFR3 aberrations (FGFR3-TACC3 and cysteine mutants) induce resistance to single agent dasatinib while others (N542K and K652E) do not. Finally, this study did not investigate the relevance of Src as a bypass compensatory pathway in the paradigm of acquired resistance to selective FGFR inhibitors. Future work will seek to determine if blockade of the Src pathway is able to re-sensitise cells in this acquired resistance context and achieve durable drug responses

In summary, we show that consistent with the clinical experience, cells bearing a subset of FGFR3 molecular alterations are intrinsically resistant to selective FGFR inhibitors. Expression of FGFR3-TACC3 and FGFR3 cysteine point mutants led to constitutive Src activation which can be exploited for therapy with the Src inhibitor dasatinib. Treatment with dasatinib in combination with selective FGFR inhibitors was highly effective in overcoming intrinsic drug resistance in urothelial cancer cells via the dual blockade of the Src and Erk1/2 pathways. This study provides proof-of-principle evidence that supports co-targeting Src and FGFR3 as a therapeutic approach to tackle drug resistance in the treatment of cancers with FGFR3 molecular alterations.

## Methods and Materials

### Cell lines

NIH-3T3 cells stably expressing WT FGFR3, FGFR3-TACC3, S249C, N542K and K652E) have previously been described (17, 20). Urothelial cancer cell lines (BFTC905, RT112M, 639V) and were kindly provided by Nicholas Turner. The remaining NIH-3T3 FGFR3 mutant expressing cell lines were generated in-house. BFTC905 and 639V cell lines were maintained in DMEM and RT112M maintained in RPMI in 10% FBS (Life Technologies) and 2mmol/L L -glutamine. NIH-3T3 cell lines were cultured with DMEM containing 10% FBS. All cell lines stably transducing expression constructs where supplemented with 100µg/mL of hygromycin (Invivogen), except for FGFR3-TACC3 which was supplemented with 800µg/mL of geneticin (Sigma-Aldrich).

### Generation of plasmids and stable cell lines

FGFR3 point mutations (S249C, S249A, R248C and R248G) were generated using a retroviral expression vector containing full-length human FGFR3 (46) by site-directed mutagenesis (SDM). SDM was performed utilising either the QuikChange Lightning SDM Kit (Agilent) and the Q5 SDM kit (New England Biolabs) following the manufacturer’s instructions (Supplementary methods). Mutations were confirmed by Sanger Sequencing (Eurofins Genomics). Avian Src constructs were obtained from Addgene and include pBabe-hygro empty vector (1765, Addgene), wildtype Src (26983, Addgene) and gatekeeper mutant Src (SRC-T338I) (26980, Addgene). Cell lines generated in-house were produced by retroviral transduction. Retroviral particles were produced in HEK293T cell line utilising 15μg of the insert plasmid, 15μg of pUMVC (8449, Addgene) and 1.88μg of pCMV-VSV-G (8454, Addgene) utilising polyethylenimine (PEI) (1mg/mL, Sigma-Aldrich) as transfection reagent. After 4 days of transfection, medium with retrovirus was harvested. NIH-3T3, BFTC905, RT112M or 639V cell lines were then transduced with the harvested retroviral medium diluted 1:4 with growth medium together with 8 μg/mL of polybrene (Sigma-Aldrich). Cells were then selected for 2 weeks with 200μg/mL of hygromycin.

### Western blotting and antibodies

Cells were grown for 48h in culture and lysed or treated with the indicated inhibitors prior to lysis in RIPA lysis buffer (50 mM Tris-HCl, pH 7.6, 150 mM NaCl, 1 % (v/v) NP40, 0.5 % (w/v) sodium deoxycholate, 0.1 % (w/v) sodium dodecyl sulphate (SDS)) with 1x Halt™ Protease and Phosphatase Inhibitor Cocktail with EDTA (Pierce, Thermo Fisher Scientific). Protein samples were prepared in 4x NuPAGE LDS Sample Buffer (Thermo Fisher Scientific) and 10x NuPAGE Sample Reducing Agent (Thermo Fisher Scientific) and boiled at 95 °C for 5 minutes. When analysing non-reducing samples, the sample reducing agent was not added to the preparation and the samples were not boiled, but the subsequent steps of the protocol remained the same. Proteins were resolved on a 4-12% gradient gel (Novex, Thermo Fisher Scientific), and transferred onto polyvinylidene fluoride (PVDF) membranes by dry transfer. Membranes were blocked and incubated with the primary and secondary HRP-conjugated antibodies in a solution with 5% (w/v) bovine serum albumin (BSA) (Sigma-Aldrich) in Tris-buffered saline solution with 0.01% (v/v) Tween (TBS-T). Antibodies used were: FGFR3 (sc-13121) 1:500 (Santa Cruz Biotechnology); FGFR3 (4574) 1:1000, phospho-Y1000 (8954) 1:2000, phospho-Src Y416 (2101) 1:500, Src (2108) 1:1000, phospho-Akt S473 (4058) 1:250-1:500, Akt (4691) 1:1000-1:2000, phospho-Erk1/2 T202/Y404 (4370) 1:2000 and Erk1/1 (4695) 1:2000-1:5000 (all from Cell Signaling); and Tubulin (T5168) 1:10,000 (Sigma-Aldrich). Anti-rabbit HRP (7074) 1:2000-1:10,000 (Cell Signaling) and anti-mouse HRP (G32-62G-1000) 1:10,000 (SignalChem). Membranes were developed with Supersignal™ West Pico PLUS Chemiluminescent Substrate (Pierce, Thermo Fisher Scientific).

### Anchorage independent cell growth in soft agar

To evaluate growth in soft agar, cells were seeded at a density of 20,000 cells/well (in 6-well plates), in a solution containing 0.35% (w/v) Noble Agar (Sigma-Aldrich) in DMEM supplemented with 10% FBS. This was layered over a 0.3% (w/v) agar solution in DMEM also supplemented with 10% FBS. Plates were incubated for 3 weeks with regular media change. Colonies were stained with Giemsa Stain Modified Solution (48900, Sigma-Aldrich) with 1 part Giemsa to 5 parts glycerol:methanol (5:24 parts). Colonies with a diameter greater than 10 pixels were analysed on ImageJ software.

### Colony formation assay

Cell growth in the presence of selected inhibitors was evaluated by colony formation assays, were cells were seeded at a density of 1000 cells/well (in 6-well plates) for 24 h, followed by addition of relevant drug treatment at the indicated concentrations. Fresh medium with drug was added twice a week for 14 days. Colonies were fixed with Carnoy’s solution (1 acetic acid: 3 methanol) and stained with 1% (w/v) crystal violet (Sigma-Aldrich). The percentage of colonies covering the plates were calculated with ImageJ software.

### Cell viability assay and small molecule inhibitor screen

NIH-3T3 cells were seeded at a density of 1000 cells/well and the urothelial cancer cells lines were seeded at a density of 3000 cells/well in a 96-well plate format. After 24h in culture, cells were treated with the selected inhibitors at the indicated concentrations (5nM up to 50µM) for a further 72h. Cell viability was assessed with the CellTitre-Glo (CTG) (Promega) reagent, as per the manufacturer’s guidelines. After evaluation of cell viability by luminescence, values were normalised against the DMSO vehicle control and plotted with a four-parameter non-linear regression on GraphPad Prism 8 (GraphPad Software Inc), where the half maximal inhibitory concentration (IC_50_) was calculated. In addition, to establish synergy in combination arms, the combination index (CI) was calculated according to the Chou-Talalay method (47).Cell viability for the small molecule inhibitor screen was also assessed with the CTG assay utilising a pre-made inhibitor library plate prepared in-house. Small molecule chemical compounds utilised: AZD4547 (A-1088), BEZ235 (N-4288), NVP-AUY922 (N-5300), Binimetinib (B-2332), Bosutinib (B-1788), Pazopanib (P-6706), Cediranib (C-4300), Crizotinib (C-7900), Ponatinib (P-7022), Dasatinib (D-3307), Foretinib (F-4185), Gefitinib (G-4408), Sorafenib (S-8599), Imatinib (I-5577), Sunitinib (S-8877), Trametinib (T-8123), Lenvatinib (L-5400), Vandetanib (V-9402), all purchased from LC Labs. MK-8776 (S2735), AZD5363 (S8019), Momelotinib (S2219), BGJ398 (S2183), NVP-TAE684 (S1108), Palbociclib (S1116), PD173074 (S1264), Ceritinib (S7083), PF562271 (S2890), Dabrafenib (S2807), GSK126 (S7061), Saracatinib (S1006), Silmitasertib (S2248), JQ1 (S7110), MK2206 (S1078), all purchased from Selleckchem. MRT67307 (SML0702), BX-795 (SML0694), both purchased from Sigma-Aldrich.

### Statistical Analysis

Statistical analysis was performed on GraphPad Prism 8 statistical software with one-way or two-way ANOVA with multiple comparisons tests. Data are presented as mean ± standard deviation (SD). The threshold for statistical significance was set to * p < 0.05, ** p < 0.01, or *** p < 0.001.

## Acknowledgements

We would like to thank Margaret Knowles for providing the FGFR3 constructs used in this study. P.H.H is supported by grants from the Institute of Cancer Research (ICR), Rosetrees Trust and Cancer Research UK (C36478/A19281) and M.K is supported by grants from Cancer Research UK (A16567).

## References

1. Di Stefano AL, Fucci A, Frattini V, Labussiere M, Mokhtari K, Zoppoli P, et al. Detection, Characterization, and Inhibition of FGFR-TACC Fusions in IDH Wild-type Glioma. Clin Cancer Res. 2015;21(14):3307–17.

2. Helsten T, Elkin S, Arthur E, Tomson BN, Carter J, Kurzrock R. The FGFR Landscape in Cancer: Analysis of 4,853 Tumors by Next-Generation Sequencing. Clin Cancer Res. 2016;22(1):259–67.

3. Robertson AG, Kim J, Al-Ahmadie H, Bellmunt J, Guo G, Cherniack AD, et al. Comprehensive Molecular Characterization of Muscle-Invasive Bladder Cancer. Cell. 2017;171(3):540–56 e25.

4. Singh D, Chan JM, Zoppoli P, Niola F, Sullivan R, Castano A, et al. Transforming fusions of FGFR and TACC genes in human glioblastoma. Science. 2012;337(6099):1231–5.

5. Cancer Genome Atlas Research N. Comprehensive molecular characterization of urothelial bladder carcinoma. Nature. 2014;507(7492):315–22.

6. Hurst CD, Platt FM, Taylor CF, Knowles MA. Novel tumor subgroups of urothelial carcinoma of the bladder defined by integrated genomic analysis. Clin Cancer Res. 2012;18(21):5865–77.

7. Robertson AG, Kim J, Al-Ahmadie H, Bellmunt J, Guo G, Cherniack AD, et al. Comprehensive Molecular Characterization of Muscle-Invasive Bladder Cancer. Cell. 2018;174(4):1033.

8. Hedegaard J, Lamy P, Nordentoft I, Algaba F, Hoyer S, Ulhoi BP, et al. Comprehensive Transcriptional Analysis of Early-Stage Urothelial Carcinoma. Cancer Cell. 2016;30(1):27–42.

9. Adar R, Monsonego-Ornan E, David P, Yayon A. Differential activation of cysteine-substitution mutants of fibroblast growth factor receptor 3 is determined by cysteine localization. J Bone Miner Res. 2002;17(5):860–8.

10. d’Avis PY, Robertson SC, Meyer AN, Bardwell WM, Webster MK, Donoghue DJ. Constitutive activation of fibroblast growth factor receptor 3 by mutations responsible for the lethal skeletal dysplasia thanatophoric dysplasia type I. Cell Growth Differ. 1998;9(1):71–8.

11. Del Piccolo N, Placone J, Hristova K. Effect of thanatophoric dysplasia type I mutations on FGFR3 dimerization. Biophys J. 2015;108(2):272–8.

12. Nelson KN, Meyer AN, Siari A, Campos AR, Motamedchaboki K, Donoghue DJ. Oncogenic Gene Fusion FGFR3-TACC3 Is Regulated by Tyrosine Phosphorylation. Mol Cancer Res. 2016;14(5):458–69.

13. Parker BC, Annala MJ, Cogdell DE, Granberg KJ, Sun Y, Ji P, et al. The tumorigenic FGFR3-TACC3 gene fusion escapes miR-99a regulation in glioblastoma. J Clin Invest. 2013;123(2):855–65.

14. Pal SK, Rosenberg JE, Hoffman-Censits JH, Berger R, Quinn DI, Galsky MD, et al. Efficacy of BGJ398, a Fibroblast Growth Factor Receptor 1-3 Inhibitor, in Patients with Previously Treated Advanced Urothelial Carcinoma with FGFR3 Alterations. Cancer Discov. 2018;8(7):812–21.

15. Loriot Y, Necchi A, Park SH, Garcia-Donas J, Huddart R, Burgess E, et al. Erdafitinib in Locally Advanced or Metastatic Urothelial Carcinoma. N Engl J Med. 2019;381(4):338–48.

16. Bellus GA, McIntosh I, Smith EA, Aylsworth AS, Kaitila I, Horton WA, et al. A recurrent mutation in the tyrosine kinase domain of fibroblast growth factor receptor 3 causes hypochondroplasia. Nat Genet. 1995;10(3):357–9.

17. Patani H, Bunney TD, Thiyagarajan N, Norman RA, Ogg D, Breed J, et al. Landscape of activating cancer mutations in FGFR kinases and their differential responses to inhibitors in clinical use. Oncotarget. 2016;7(17):24252–68.

18. di Martino E, L’Hote CG, Kennedy W, Tomlinson DC, Knowles MA. Mutant fibroblast growth factor receptor 3 induces intracellular signaling and cellular transformation in a cell type- and mutation-specific manner. Oncogene. 2009;28(48):4306–16.

19. Williams SV, Hurst CD, Knowles MA. Oncogenic FGFR3 gene fusions in bladder cancer. Hum Mol Genet. 2013;22(4):795–803.

20. Bunney TD, Wan S, Thiyagarajan N, Sutto L, Williams SV, Ashford P, et al. The Effect of Mutations on Drug Sensitivity and Kinase Activity of Fibroblast Growth Factor Receptors: A Combined Experimental and Theoretical Study. EBioMedicine. 2015;2(3):194–204.

21. Chell V, Balmanno K, Little AS, Wilson M, Andrews S, Blockley L, et al. Tumour cell responses to new fibroblast growth factor receptor tyrosine kinase inhibitors and identification of a gatekeeper mutation in FGFR3 as a mechanism of acquired resistance. Oncogene. 2013;32(25):3059–70.

22. Guagnano V, Kauffmann A, Wohrle S, Stamm C, Ito M, Barys L, et al. FGFR genetic alterations predict for sensitivity to NVP-BGJ398, a selective pan-FGFR inhibitor. Cancer Discov. 2012;2(12):1118–33.

23. Herrera-Abreu MT, Pearson A, Campbell J, Shnyder SD, Knowles MA, Ashworth A, et al. Parallel RNA interference screens identify EGFR activation as an escape mechanism in FGFR3-mutant cancer. Cancer Discov. 2013;3(9):1058–71.

24. Guagnano V, Furet P, Spanka C, Bordas V, Le Douget M, Stamm C, et al. Discovery of 3-(2,6-dichloro-3,5-dimethoxy-phenyl)-1-{6-[4-(4-ethyl-piperazin-1-yl)-phenylamin o]-pyrimidin-4-yl}-1-methyl-urea (NVP-BGJ398), a potent and selective inhibitor of the fibroblast growth factor receptor family of receptor tyrosine kinase. J Med Chem. 2011;54(20):7066–83.

25. Kwarcinski FE, Brandvold KR, Phadke S, Beleh OM, Johnson TK, Meagher JL, et al. Conformation-Selective Analogues of Dasatinib Reveal Insight into Kinase Inhibitor Binding and Selectivity. ACS Chem Biol. 2016;11(5):1296–304.

26. Li J, Rix U, Fang B, Bai Y, Edwards A, Colinge J, et al. A chemical and phosphoproteomic characterization of dasatinib action in lung cancer. Nat Chem Biol. 2010;6(4):291–9.

27. Lopez JS, Banerji U. Combine and conquer: challenges for targeted therapy combinations in early phase trials. Nat Rev Clin Oncol. 2017;14(1):57–66.

28. Tan AC, Vyse S, Huang PH. Exploiting receptor tyrosine kinase co-activation for cancer therapy. Drug Discov Today. 2017;22(1):72–84.

29. Nakanishi Y, Akiyama N, Tsukaguchi T, Fujii T, Satoh Y, Ishii N, et al. Mechanism of Oncogenic Signal Activation by the Novel Fusion Kinase FGFR3-BAIAP2L1. Mol Cancer Ther. 2015;14(3):704–12.

30. Acquaviva J, He S, Zhang C, Jimenez JP, Nagai M, Sang J, et al. FGFR3 translocations in bladder cancer: differential sensitivity to HSP90 inhibition based on drug metabolism. Mol Cancer Res. 2014;12(7):1042–54.

31. Elliott AY, Bronson DL, Cervenka J, Stein N, Fraley EE. Properties of cell lines established from transitional cell cancers of the human urinary tract. Cancer Res. 1977;37(5):1279–89.

32. Suwaki N, Vanhecke E, Atkins KM, Graf M, Swabey K, Huang P, et al. A HIF-regulated VHL-PTP1B-Src signaling axis identifies a therapeutic target in renal cell carcinoma. Sci Transl Med. 2011;3(85):85ra47.

33. Goyal L, Saha SK, Liu LY, Siravegna G, Leshchiner I, Ahronian LG, et al. Polyclonal Secondary FGFR2 Mutations Drive Acquired Resistance to FGFR Inhibition in Patients with FGFR2 Fusion-Positive Cholangiocarcinoma. Cancer Discov. 2017;7(3):252–63.

34. Goyal L, Shi L, Liu LY, Fece de la Cruz F, Lennerz JK, Raghavan S, et al. TAS-120 Overcomes Resistance to ATP-Competitive FGFR Inhibitors in Patients with FGFR2 Fusion-Positive Intrahepatic Cholangiocarcinoma. Cancer Discov. 2019;9(8):1064–79.

35. Datta J, Damodaran S, Parks H, Ocrainiciuc C, Miya J, Yu L, et al. Akt Activation Mediates Acquired Resistance to Fibroblast Growth Factor Receptor Inhibitor BGJ398. Mol Cancer Ther. 2017;16(4):614–24.

36. Wang J, Mikse O, Liao RG, Li Y, Tan L, Janne PA, et al. Ligand-associated ERBB2/3 activation confers acquired resistance to FGFR inhibition in FGFR3-dependent cancer cells. Oncogene. 2015;34(17):2167–77.

37. Levitt JM, Yamashita H, Jian W, Lerner SP, Sonpavde G. Dasatinib is preclinically active against Src-overexpressing human transitional cell carcinoma of the urothelium with activated Src signaling. Mol Cancer Ther. 2010;9(5):1128–35.

38. Thomas S, Overdevest JB, Nitz MD, Williams PD, Owens CR, Sanchez-Carbayo M, et al. Src and caveolin-1 reciprocally regulate metastasis via a common downstream signaling pathway in bladder cancer. Cancer Res. 2011;71(3):832–41.

39. Hahn NM, Knudsen BS, Daneshmand S, Koch MO, Bihrle R, Foster RS, et al. Neoadjuvant dasatinib for muscle-invasive bladder cancer with tissue analysis of biologic activity. Urol Oncol. 2016;34(1):4 e11–7.

40. Xu AM, Huang PH. Receptor tyrosine kinase coactivation networks in cancer. Cancer Res. 2010;70(10):3857–60.

41. Song L, Morris M, Bagui T, Lee FY, Jove R, Haura EB. Dasatinib (BMS-354825) selectively induces apoptosis in lung cancer cells dependent on epidermal growth factor receptor signaling for survival. Cancer Res. 2006;66(11):5542–8.

42. Zhang J, Kalyankrishna S, Wislez M, Thilaganathan N, Saigal B, Wei W, et al. SRC-family kinases are activated in non-small cell lung cancer and promote the survival of epidermal growth factor receptor-dependent cell lines. Am J Pathol. 2007;170(1):366–76.

43. Haura EB, Tanvetyanon T, Chiappori A, Williams C, Simon G, Antonia S, et al. Phase I/II study of the Src inhibitor dasatinib in combination with erlotinib in advanced non-small-cell lung cancer. J Clin Oncol. 2010;28(8):1387–94.

44. Dhawan A, Nichol D, Kinose F, Abazeed ME, Marusyk A, Haura EB, et al. Collateral sensitivity networks reveal evolutionary instability and novel treatment strategies in ALK mutated non-small cell lung cancer. Sci Rep. 2017;7(1):1232.

45. Zhao B, Sedlak JC, Srinivas R, Creixell P, Pritchard JR, Tidor B, et al. Exploiting Temporal Collateral Sensitivity in Tumor Clonal Evolution. Cell. 2016;165(1):234–46.

46. Tomlinson DC, L’Hote CG, Kennedy W, Pitt E, Knowles MA. Alternative splicing of fibroblast growth factor receptor 3 produces a secreted isoform that inhibits fibroblast growth factor-induced proliferation and is repressed in urothelial carcinoma cell lines. Cancer Res. 2005;65(22):10441–9.

47. Chou TC. Drug combination studies and their synergy quantification using the Chou-Talalay method. Cancer Res. 2010;70(2):440–6.

