## Supplementary figures and images for "Targeting the Src pathway enhances the efficacy of selective FGFR inhibitors in cancers with FGFR3 alterations"

### Supplemental Figures

Supplemental Figure 1

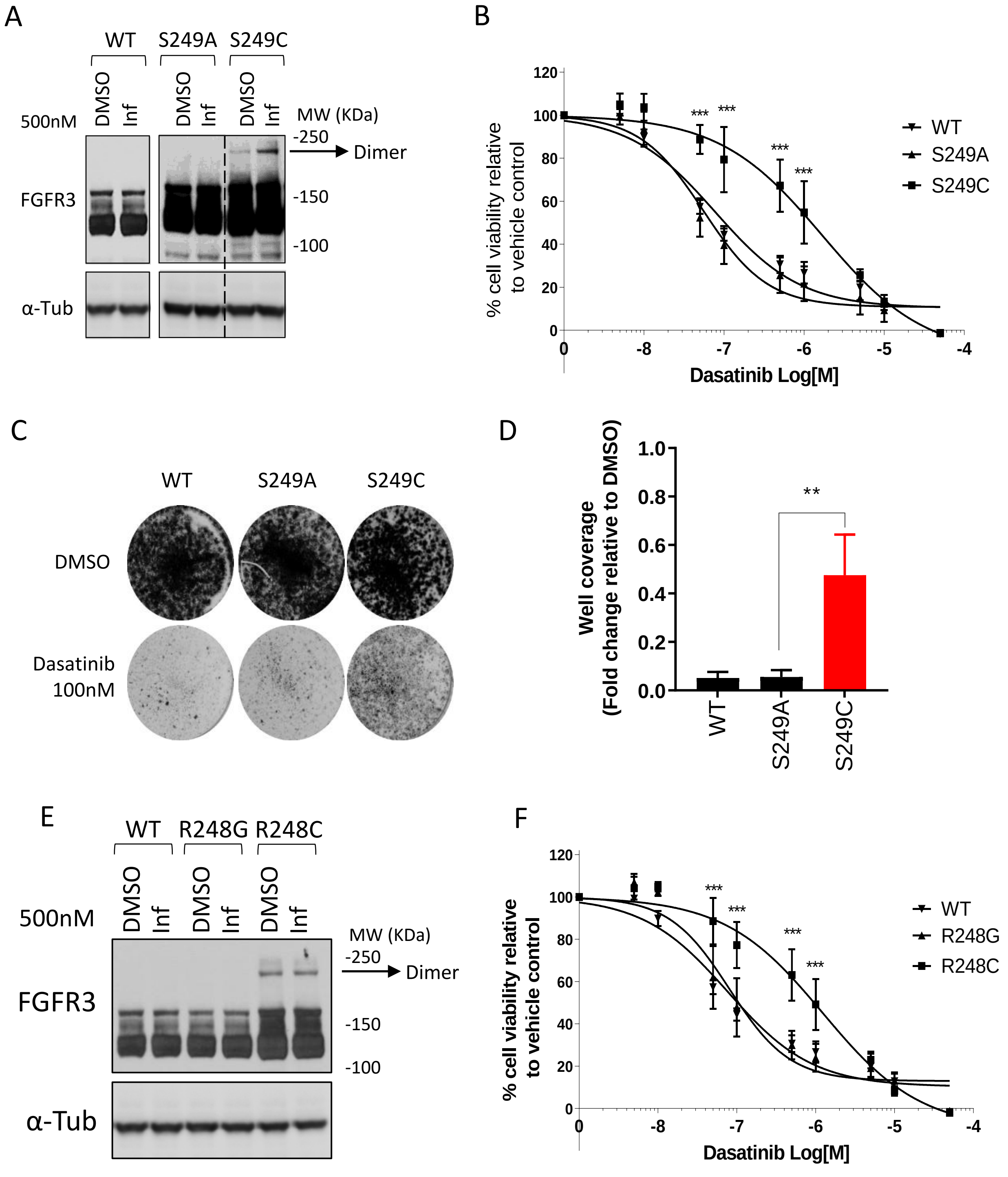

Supplemental Figure 2

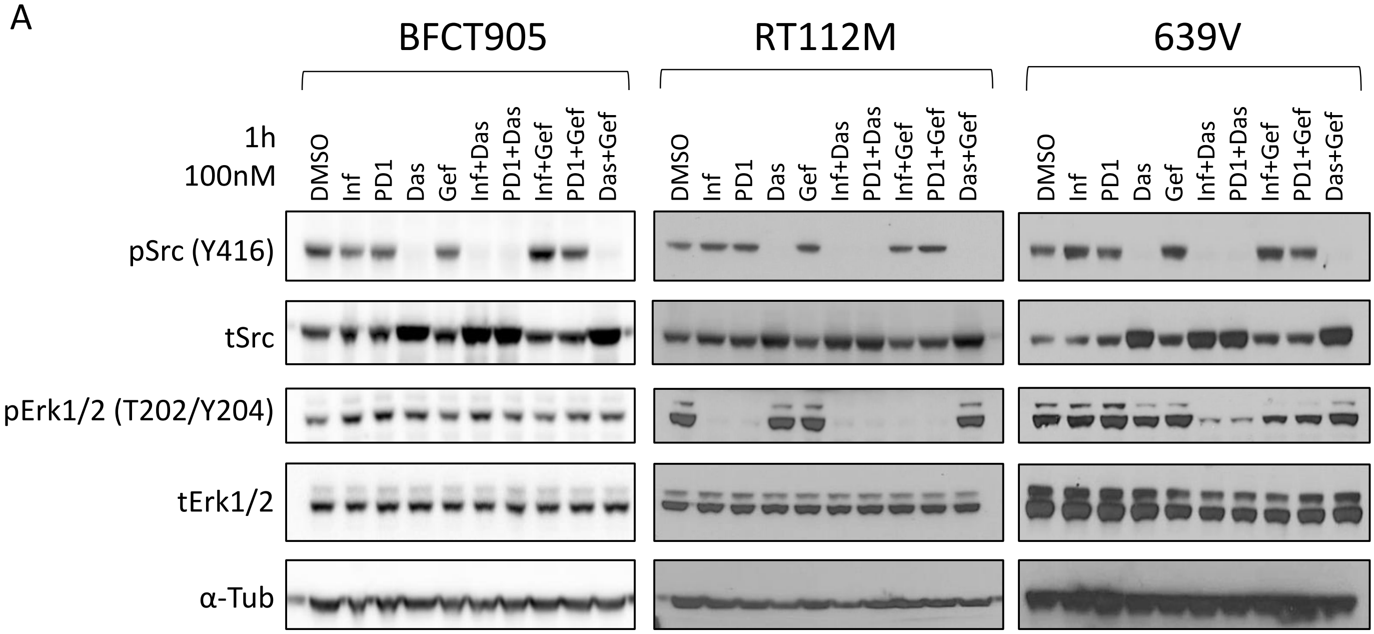


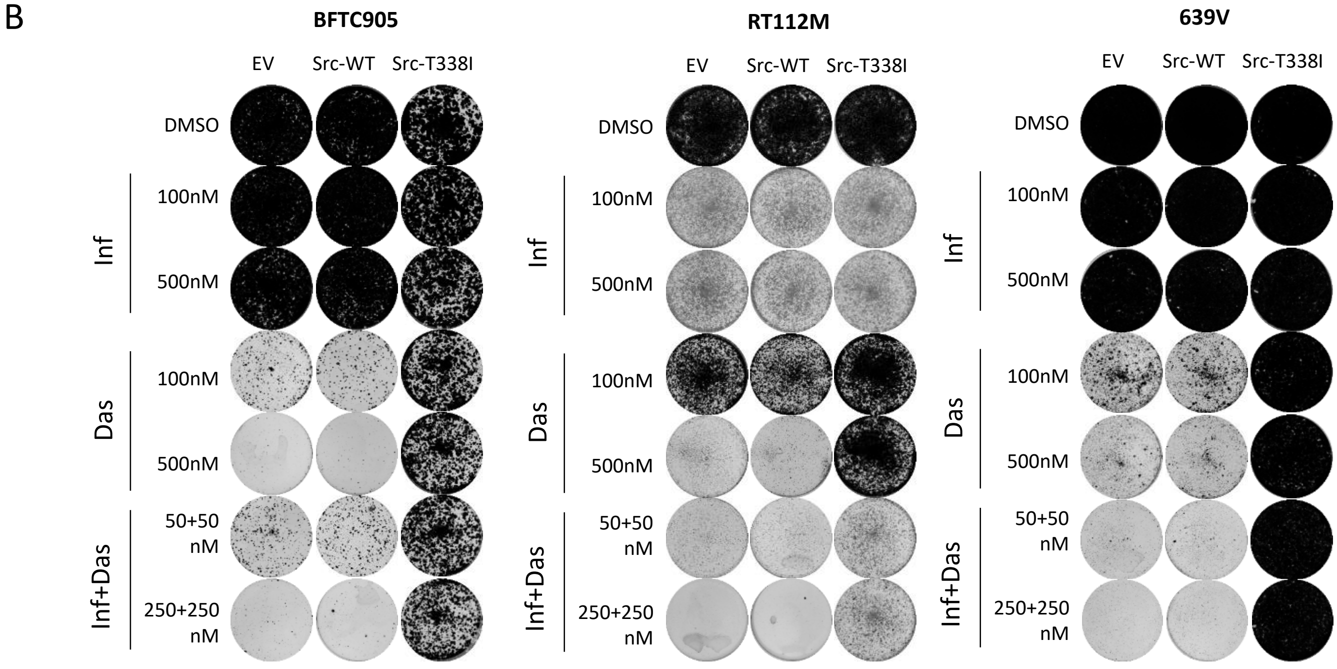
