## Supplemental methods for "Targeting the Src pathway enhances the efficacy of selective FGFR inhibitors in cancers with FGFR3 alterations"

List of primer sequences used to generate FGF3 mutations in the WT template pFB-Hyg-FGFR3.

*Primers were originally designed to generate the mutant R248A (targeted sequence AGGCC), but Sanger sequencing results showed that the generated mutant corresponded to R248G (sequence AGCGC).

| **FGFR3 mutation** | | **Primers (5' → 3')** | |
| --- | --- | --- | --- |
| **Used with QuikChange Lightning SDM Kit** | | | |
| R248C | CGC → TGC | Forward | GACGTGCTGGAG**T**GCTCCCCGCACC |
|  |  | Reverse | GGTGCGGGGAGC**A**CTCCAGCACGTC |
| S249C | TCC → TGC | Forward | ACGTGCTGGAGCGCT**G**CCCGCACC |
|  |  | Reverse | GGTGCGGG**C**AGCGCTCCAGCACGT |
| **Used with Q5 SDM kit** | | | |
| R248A* | CGC → GCC | Forward | CGTGCTGGAG**GC**CTCCCCGCAC |
|  |  | Reverse | TCCAGCGTGTACGTCTGC |
| S249A | TCC → GCC | Forward | GCTGGAGCGC**G**CCCCGCACCGGC |
|  |  | Reverse | ACGTCCAGCGTGTACGTCTGCCGGATG |
