## Supplemental Tables for "Targeting the Src pathway enhances the efficacy of selective FGFR inhibitors in cancers with FGFR3 alterations"

Supplemental table 1 – List of inhibitors utilised for the targeted small molecule inhibitor screen. The major cellular targets of inhibitors included in the screen are indicated.

| **Compound** | **Target** | **Major pathway** |
| --- | --- | --- |
| AZD4547 | FGFR1-3 | FGFR |
| AZD5363 | Akt | PI3K/Akt/mTOR |
| BEZ235 | PI3K, mTOR | PI3K/Akt/mTOR |
| BGJ398 | FGFR1-3 | FGFR |
| Binimetinib | MEK1/2 | MAPK |
| Bosutinib | Src | Src |
| BX-795 | PDK1, TBK1, IKKϵ | NFkB and TBK1 |
| Cediranib | Broad-spectrum: VEGFR, Flt1/4, KIT | Broad-spectrum TKi |
| Ceritinib | ALK | ALK |
| Crizotinib | Met, ALK | ALK |
| Dabrafenib | BRAF V600E | MAPK |
| Dasatinib | Broad-spectrum: SFK, Abl, KIT | Src |
| Foretinib | Broad-spectrum: VEGFR, MET | Broad-spectrum TKi |
| Imatinib | Broad-spectrum: Abl, KIT, PDGFR | Broad-spectrum TKi |
| JQ1 | BRD1-4 | BET bromodomain family |
| Lenvatinib | Broad-spectrum: VEGFR2/3 | Broad-spectrum TKi |
| MK2206 | Akt | PI3K/Akt/mTOR |
| MK-8776 | CHK1 | ATR-CHK1 |
| Momelotinib | JAK1/2 | JAK/STAT |
| MRT67307 | IKKϵ, TBK1 | NFkB and TBK1 |
| NVP-AUY922 | Hsp90 | Hsp90 |
| NVP-TAE684 | ALK | ALK |
| Palbociclib | CDK4, CDK6 | Cell cycle regulator |
| Pazopanib | Broad-spectrum: VEGFR1-3, PDGFR | Broad-spectrum TKi |
| PF562271 | FAK, Pyk2 | FAK |
| Ponatinib | Abl, PDGFRα, VEGFR2, FGFR1, Src | Broad-spectrum TKi |
| GSK126 | EZH2 | EZH2 methyltransferase |
| Silmitasertib | CK2 | CK2 |
| Sorafenib | Broad-spectrum: VEGFR2, BRAF, RAF | Broad-spectrum TKi |
| Sunitinib | Broad-spectrum: VEGFR2, PDGFRß | Broad-spectrum TKi |
| Trametinib | MEK1/2 | MAPK |
| Vandetanib | Broad-spectrum: VEGFR2/3, EGFR | VEGFR |
