## Supplemental figure legends for "Targeting the Src pathway enhances the efficacy of selective FGFR inhibitors in cancers with FGFR3 alterations"

**Supplemental Figure 1.** (A) Immunoblot of FGFR3 dimers and monomers in NIH-3T3 cells bearing WT FGFR3, S249A and S249C mutants treated with 500nM of infigratinib (Inf) or DMSO control for 1 hour under non-reducing native PAGE conditions. Molecular weight of FGFR3 dimers are indicated by the arrow. (B) Dose-response curves of the WT FGFR3, S249A and S249C cell lines upon treatment with dasatinib for 72 hours. Cell viability is normalised to DMSO control treatment (n=3). Statistical analysis of S249C mutant cell line versus S249A mutant was performed by two-way ANOVA with Dunnett’s post-hoc multiple comparison adjusted p-value at each drug dose (***p<0.001). (C) Representative images of long term colony formation assay in the WT FGFR3, S249A and S249C cell lines upon treatment with dasatinib or DMSO control at the indicated dose for 2 weeks. (D) Bar plots showing the quantification of well coverage of the colony formation assay in panel I. Data for each cell line is normalised to DMSO control treatment (n=3). Statistical analysis of S249C versus S249A was performed by paired Student’s t test (**p<0.01). (E) Immunoblot of FGFR3 dimers and monomers in NIH-3T3 cells bearing WT FGFR3, R248G and R248C mutants treated with 500nM of infigratinib (Inf) or DMSO control for 1 hour under non-reducing native PAGE conditions. Molecular weight of FGFR3 dimers are indicated by the arrow. (F) Dose-response curves of the WT FGFR3, R249G and R249C cell lines upon treatment with dasatinib for 72 hours. Cell viability is normalised to DMSO control treatment (n=3). Statistical analysis of R249C mutant cell line versus R249G mutant was performed by two-way ANOVA with Dunnett’s post-hoc multiple comparison adjusted p-value at each drug dose (***p<0.001). Data presented for (B), (D) and (F) represent mean ± SD.

**Supplemental Figure 2.** (A) Immunoblot of Erk1/2 and Src phosphorylation levels in urothelial cancer cells treated with single agent selective FGFR inhibitor, dasatinib, gefitinib and in combination for 1 hour. The combination arm was administered in a 1:1 ratio, with the indicated concentration representing the total drug concentration used. Inf – infigratinib, PD – PD173074, Gef – gefitinib, Das – dasatinib. (B) Representative images of long term colony formation assay in urothelial cancer cell line panel that have been transduced with empty vector (EV), wildtype (WT) Src or gatekeeper mutant Src (SRC-T338I) and treated with infigratinib, dasatinib as single agent or in combination as well as DMSO control at the indicated doses for 2 weeks.
